# Development of a novel allograft model of prostate cancer: a new tool to inform clinical translation

**DOI:** 10.1101/2020.02.16.942789

**Authors:** Charles M. Haughey, Debayan Mukherjee, Rebecca E. Steele, Amy Popple, Lara Dura-Perez, Adam Pickard, Suneil Jain, Paul B. Mullan, Rich Williams, Pedro Oliveira, Niamh E. Buckley, Jamie Honeychurch, Simon McDade, Timothy Illidge, Ian G. Mills, Sharon L. Eddie

**Affiliations:** Centre for Cancer Research and Cell Biology, Queen’s University, Belfast, UK; Targeted Therapy Group, Division of Cancer Sciences, Faculty of Biology Medicine and Health, The University of Manchester, UK; Wellcome Centre for Cell Matrix Research, University of Manchester, M13 9PL, UK; Nuffield Department of Surgical Sciences, John Radcliffe Hospital, University of Oxford, Oxford, United Kingdom; Department of Pathology, the Christie Hospital Foundation Trust, Manchester, UK

## Abstract

**Background:** Allograft models enable characterisation of genomic drivers and treatment responses by modelling immune and micro-environmental changes more accurately than xenografts. Despite this, few models are available to the prostate cancer (PCa) community. This study presents a novel allograft model of high-risk, localised PCa. In characterising this model we have focused on its response to radiotherapy (RT).

**Methods:** The DVL3 cell line was derived from a transgenic trp53^-/-^/Pten^-/-^ mouse model and characterised both *in vitro* and *in vivo*. The DVL3 cells were allografted and response to RT was investigated and compared to the TRAMP-C1 model. Extensive tumour profiling in the DVL3 model was performed using flow cytometry, immunohistochemistry and RNA-seq.

**Results:** *In vitro* the DVL3 cells expressed basal and luminal markers. DVL3 cells formed tumours with distinct glandular morphology which expressed androgen receptor, similar to human localised PC. DVL3 tumour growth was delayed following administration of fractionated RT, with infiltration of myeloid derived suppressor cells (MDSC).

**Conclusions:** The DVL3 allograft model represents substantial progress in PCa modelling, displaying; luminal differentiation, strong AR expression, and an immunosuppressive microenvironment, similar to observations in high-risk PCa patients. This model is ideally suited for development and validation of novel therapeutics, in particular immune-modulatory agents in combination with RT.

## Background

Prostate cancer (PCa) is the most common cancer in men and the fifth leading cause of cancer deaths ^1^. It is estimated that 1.3 million new cases of PCa were diagnosed worldwide in 2018 alone ^1^. The vast majority of PCa (91%) is localised disease at diagnosis ^2^, and can be treated with a range of therapeutic modalities including surgery, androgen-deprivation therapy (ADT) and radiotherapy (RT). Unfortunately, roughly 30% of high-risk localised PCa will develop into aggressive metastatic disease ^3^, with limited treatment options. The current standard of care for these patients is ADT, but this has many adverse side effects and has a median relapse rate of only 11 months ^4^.

As a result, efforts have focused on the use of RT to treat both localised and metastatic disease. RT delivers comparable patient outcomes to radical prostatectomy; survival and disease control, with less impact on patient quality of life ^5^. However, one major challenge in further enhancing RT responses, by combining this modality with other treatments, is the paucity of appropriate preclinical PCa models in which to test these combinations.

The most commonly used immune-competent model is the TRAMP-C1 murine cell line, which can be allografted subcutaneously to generate more uniform, accessible prostate tumours ^6^. The TRAMP-C1 model was generated by engineered expression of SV40 large T-antigen. Unfortunately, TRAMP-C1 allografts develop neuroendocrine tumours which are rare clinically, rather than adenocarcinomas which are most often seen in patients with localised prostate cancer (7).

Consequently, novel PCa models that more accurately recapitulate human disease are urgently required. We developed an allograft model from the trp53^-/-^/Pten^-/-^ transgenic mouse tumour; the DVL3 cell-line (derived from tumour in the **D**orsal, **V**entral and **L**ateral prostate lobes). DVL3 cells develop tumours in immune competent mice that retain morphological, lineage and immune characteristics of localised high-risk PCa. These tumours respond to RT and retain androgen receptor (AR) expression. This new model is ideal for future pre-clinical evaluation of novel treatment combinations including immune therapeutic agents.

## Materials and Methods

### Cell Line Derivation

Tissue fragments were centrifuged at 2000 rpm to remove extraneous blood and fat, PBS was removed, and remaining tissue digested in 1mg/ml collagenase/dispase in PBS for 30min at 37°C. Tissue fragments were further manually dissociated and the supernatant containing prostate cells collected. Collagnease/dispase was inactivated with EDTA at this time. The digestion process was repeated twice with remaining tissue fragments. Supernatant from the digestions was pooled and centrifuged at 2000 rpm and prostate cells were re-suspended in RPMI supplemented with 10% FBS, 1X pen/strep, 100nM DHT and rock inhibitor. These cells were then incubated for 10 minutes at 37°C to allow for prostate fibroblasts to adhere to the tissue culture plastic. Remaining cell suspension, enriched for prostate epithelium was transferred to a new culture flask. This process of differential plating, allowing for initial adherence of fibroblasts and removal and retention of less adherent epithelial cells, was repeated for 10 passages, at which time the rock inhibitor was removed and the population remained homogenous.

### Cell Line Generation and Maintenance

Mouse prostate epithelial cells (MPEC) were generated from dorsal, ventral and lateral lobes of the prostate from Probasin Cre^-/-^ (Pb-Cre4) mice. Murine prostate cancer cells (DVL3) were generated from tumours derived from the dorsal, ventral and lateral prostate lobes of a trp53^-/-^/Pten^-/-^ Pb-Cre4 mouse ^7^. Tissue was manually dissociated under sterile conditions and cell lines were generated as described in **Cell Line Derivation and Supplementary Figure-1A**.

The TRAMP-C1 murine prostate carcinoma cells were purchased from ATCC and maintained in DMEM high glucose medium, supplemented with 4mM L-Glutamine, 5% FBS, 5% Nu Serum, 0.005mg/ml of Bovine insulin, and 10nM Dehydroisoandrosterone (Sigma). The MPEC and DVL3 cell lines were maintained in RPMI-media supplemented with 10% FBS, L-Glutamine and 100nM DHT.

### RNA Isolation and cDNA synthesis

Total RNA from cells was isolated using TriPure Isolation reagent (Roche) and phenol-chloroform extraction and cDNA synthesised using first strand cDNA (Roche) according to the respective manufacturer’s instructions.

### qRT-PCR

mRNA levels were determined by RT-PCR and measured on Roche LighCycler®480 II. All primers were obtained from Eurofins and sequences are detailed in (**Supplemental Table-1**) mRNA expression was quantified relative to house-keeping genes; B-actin.

### Western blotting

Cells were lysed with Erythrocyte Lysis Buffer (0.5mM DTT, 5mM EDTA (pH 8), 50mM HEPES (pH7.5), 0.1% IGEPAL, 250mM NaCl) supplemented with Protease Inhibitor Cocktail (Roche), centrifuged and supernatant containing protein analysed by SDS Page (Invitrogen). Proteins were transferred onto a nitrocellulose membrane, blocked for 1 hour in 5% milk TBS-T prior to incubation with primary antibody at 4°C overnight, concentrations are detailed in (**Supplementary Table-2**). Complementary secondary antibodies were used at 1:3000. Membranes were visualised on a Syngene G:Box imager using reagent (Merck).

### Allograft modelling

C57BL/6 male mice (8 weeks old) were obtained from Harlan, UK. All animal experiments were performed under United Kingdom Home Office License (PPL2775; PCC943F76) held at Queen’s University Belfast or the CRUK Manchester institute, University of Manchester. Prior each *in vivo* experiment, cells were screened for mycoplasma contamination and MHV. Mice were housed on a 12/12 light/dark cycle and were given filtered water and fed ad libitum. Mice were inoculated subcutaneously with either 5×10^6^ TRAMP-C1, 1×10^6^ DVL3 cells or 1×10^6^ mPEC cells.

### Radiotherapy

Upon tumour establishment (4-6 weeks post inoculation), mice were randomised to treatment groups. Irradiation was performed when the tumours were between (∼100mm^3^). The tumour bearing mice were placed in a lead jig with an opening for the tumour and shielding for the rest of the body. Radiotherapy was delivered using an AGO cell X-Ray unit and XSTRAHL *in vivo* irradiator at a dose rate of 2.086Gy/ minute. Tumour bearing mice received either single dose of 8Gy irradiation, or 5 fractions of 2Gy delivered over 5 consecutive days. Experimental groups contained at least 4-5 mice per group.

### Sample Preparation

Tumour bearing mice were sacrificed at the indicated timepoints, tumours excised and fixed in 4% buffered formalin (Sigma Aldrich, UK) for 24 hours or collected in media for tumour disaggregation. The formalin fixed tumour samples were transferred to 70% ethanol and processed to FFPE blocks.

### Immunohistochemistry

Briefly, the slides were deparaffinised in xylene followed by rehydration in ethanol. Antigen retrieval was performed using pH 6 citrate buffer, followed by 3% H_2_0_2_ block. Slides were incubated with 10% serum, prior to incubation with primary antibody overnight at 4° (**Supplementary Table-2**). Primary antibody was detected with either HRP detection kit or biotinylated secondary antibody followed by ABC detection kit (Vector Labs, USA). Slides were briefly incubated in DAB substrate (Vector Labs, USA), washed in water, and counter-stained using haematoxylin. For multiplex staining of mouse FFPE tumour sections, the opal TSA detection system (Opal 520®, 570® and 650®) were applied to sections following manufacturer’s instruction and run on Leica BOND Rx automated system. The slides were counterstained with DAPI. Image analysis and quantification was performed using Definiens Tissue Phenomics Software.

### Immunocytochemistry

Cells were seeded onto glass slides pre-coated with collagen type II (BD Biosciences) and left to adhere for 24 hours before fixation with 4% Paraformaldehyde at RT. Cells were permeabilised with 0.1% Triton X and blocked with 3% FBS-PBS. Primary antibody was added to slides and incubated for 1 hour at RT followed by incubation in secondary antibody for 1 hour at RT (**Supplemental Table-2**). Coverslips were mounted onto glass slides using Mounting media (Invitrogen) containing Dapi and imaged.

### Flow cytometry on cell lines

Cells were incubated with primary antibody conjugated fluorophore for 30 minutes at 4°C, washed with PBS and analysed on the Introducing the BD Accuri™ C6 Plus according to the manufacturer’s instructions.

### Flow cytometry on tumour tissue

To obtain single cell suspensions, tumours were processed using a gentleMacs dissociator and a murine dissociation kit (Miltenyi Biotec). For staining of cells, non-specific binding was blocked with rat anti-CD16/CD32 Fc block on ice. Cells were incubated with Gr-1 FITC, CD11b-APC (eBiosceince), washed in 1% FCS/PBS. For analysis, live cells were gated using vital dye exclusion (Invitrogen) and population phenotyped on FACs Canto (BD Bioscience) and analysed using Flow Jo software. An example of the gating strategy employed for selection of either CD45^-^, CD45^+^, or CD45^+^CD11b^+^Gr1^+^cells is provided in **Supplementary Figure 2**.

### RNA extraction for RNA seq analysis

RNA from FFPE mouse tumour tissue was extracted. Briefly, 2-4 (10uM) sections were transferred into RNAase free microcentrifuge tube. The paraffin was removed by incubating in xylene and ethanol. For lysate and total RNA purification, digestion buffer and proteinase-K was added to the samples as per manufacturer’s instructions (Norgen kit). The samples were spun briefly followed by transferring the supernatant to a new RNAase free microcentrifuge tube. The RNA containing tubes were incubated for 15 minutes at 80°C. The lysates were then passed through RNA purification microcolumn and centrifuged for 1 minute at 14,000 RPM. The microcolumns were washed according to manufacturer’s instructions and the RNA eluted using the Elution solution (Norgen Kit).

### RNA library preparation, sequencing and analysis

RNA libraries preparations were generated using the Quant-seq 3’ mRNA-Seq FWD kit (Lexogen PART NO k15.96) according to manufacturer’s instructions, with 500ng input RNA. FFPE-derived RNA was prepared using the suggested manufacturer’s instructions. Libraries were pooled and sequenced on the NextSeq 500 using the FMHLS genomics CTU, yielding an average of 10M reads per sample. FASTQ files were aligned to the mm10 genomic reference using STAR aligner ^8^ and counts quantified at a gene level with HTseq-Counts ^9^ to yield 5-8M mapped reads per sample. Differential gene expression was performed between the irradiated and un-irradiated samples using the DESEQ 2. Differentially expressed genes were filtered; P-Value <0.05 and Log2 Fold-Change ≥1 (up-regulated) or ≤-1 (down-regulated). Pathway analysis was performed on filtered genelists using the Reactome database ^10^ in Enrichr ^11,12^. To identify enrichment of immune populations with the RT tumours vs untreated, enrichment analysis was performed using the open assess tool gene set enrichment analysis (GSEA) ^13,14^ with 1000 gene label permutations. Immune cells genes for macrophages, NK cells, T-cells and MDSC were taken from pervious published studies ^15^–^17^.

## Results

### Novel cell models express clinically relevant markers

Murine cell lines were generated via spontaneous immortalisation of normal prostate epithelium (mPECs) and prostate tumours (DVL3) **(Supplemental Figure-1A)**. Prior to *in vivo* characterisation, the cell lines were characterised *in vitro* to determine their similarity to patient disease and compared to the established TRAMP-C1 model. PCa arises from glandular epithelial cells of the prostate, and retains expression of classical prostate markers including cytokeratins 5 and 8 (CK5 and CK8). qRT-- PCR revealed significantly higher mRNA expression of CK5 and CK8 in mPEC and DVL3 cells compared to the TRAMP-C1 cells **(Figure-1A)**. Protein expression of CK8 was also notably higher in mPEC and DVL3 cells compared to TRAMP-C1 and the DVL3 cells did not express Pten **(Figure-1A and C)**. All models expressed AR at both the mRNA and protein level **(Figure-1B-C)**. Although expression of AR was higher in TRAMP-C1 cells, DVL3 cells responded to androgens and were equally responsive to Enzalutamide **(Supplemental Figure-1B)**. Interestingly, CK5 protein expression was not detectable in any of the prostate models when examined via immunocytochemistry.

**Figure 1:**
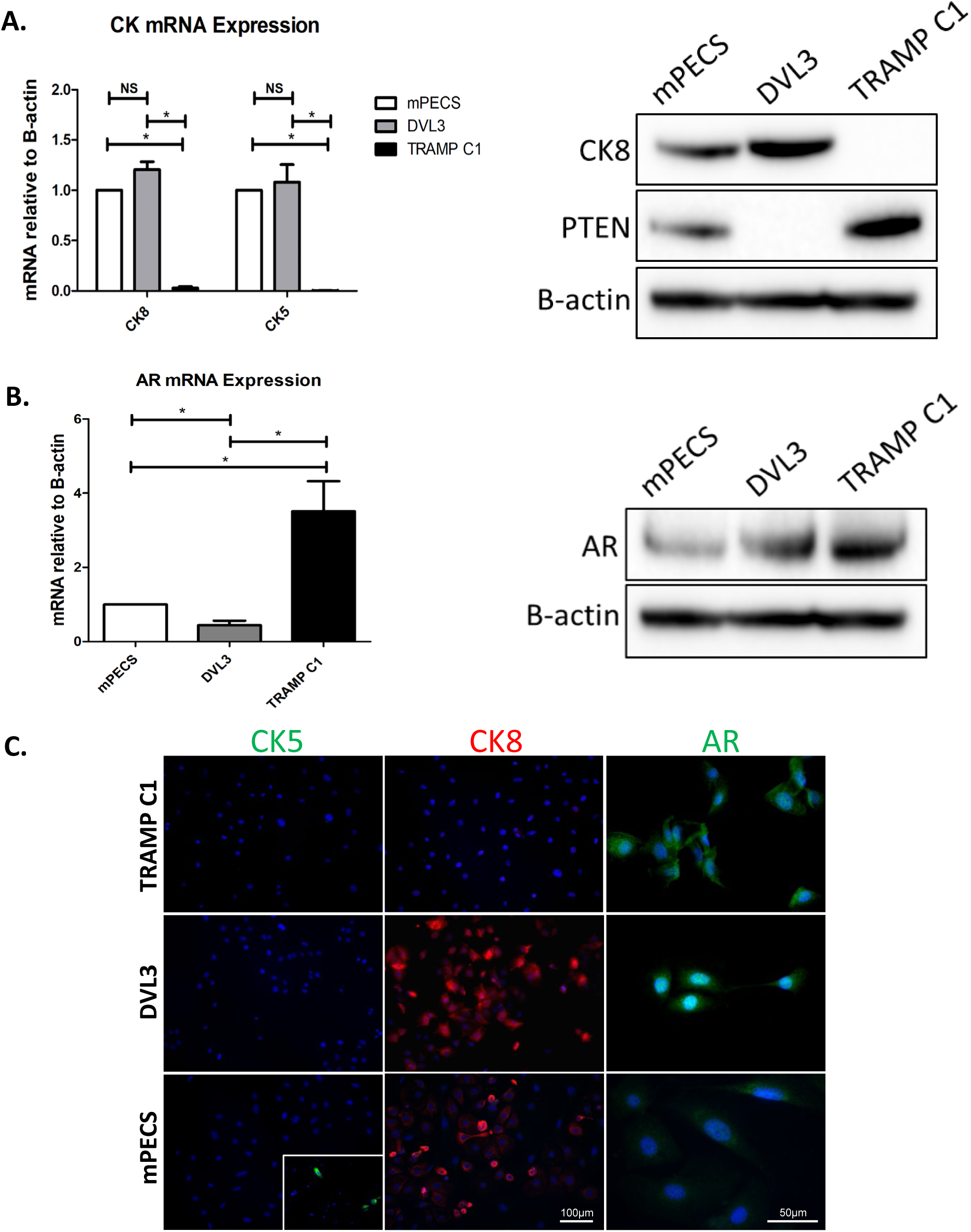
*In vitro* characterisation of mPEC and DVL3 cells demonstrates their resemblance to patient tissues. **(A)** Both novel cell models (mPEC and DVL3) express cytokeratins 5 and 8 (CK5 and CK8) mRNA at significantly higher levels than TRAMP C1 cells, as analysed via qRT-PCR. Expression of CK8 (luminal marker) was also evident at the protein level. Importantly, DVL3 cells demonstrate loss of the clinically relevant tumour suppressor, PTEN. B-actin provided as a loading control. **(B)** Androgen receptor (AR) mRNA and protein were expressed in all models examined, as assessed via qRT-PCR and western blotting, with TRAMP C1 demonstrating the highest expression of all models. **(C)** Immunocytochemical staining was consistent with qRT-PCR and western blotting results for CK8 (red) and AR (green). CK5 protein (green) was not detectable in any of the models (MCF7 cells provided as a positive control, inset). Dapi (blue) used as a counterstain. Data represents mean ± SEM of at least n=3 replicates. * denotes p≤0.05, NS denotes non-significant, as determined by unpaired t-test.

### DVL3 engraftment in immunocompetent mice forms tumours, which accurately models human adenocarcinomas

To establish tumorigenic potential, mPEC, DVL3 and TRAMP-C1 cells were subcutaneously allografted into wild-type C57BL/6 mice, as all cell lines were originally generated from the C57BL/6 strain. Mice engrafted with mPEC cells did not develop any sign of disease after 12 weeks (data not shown), consistent with their status as untransformed but spontaneously immortalised wild-type prostate epithelial cells. DVL3 tumours grew at a similar rate to the TRAMP-C1 model **(Figure-2A)**. Resultant tumours were characterised using a panel of relevant histological markers to determine their relevance as a model of human disease **(Figure-2B)**.

**Figure 2:**
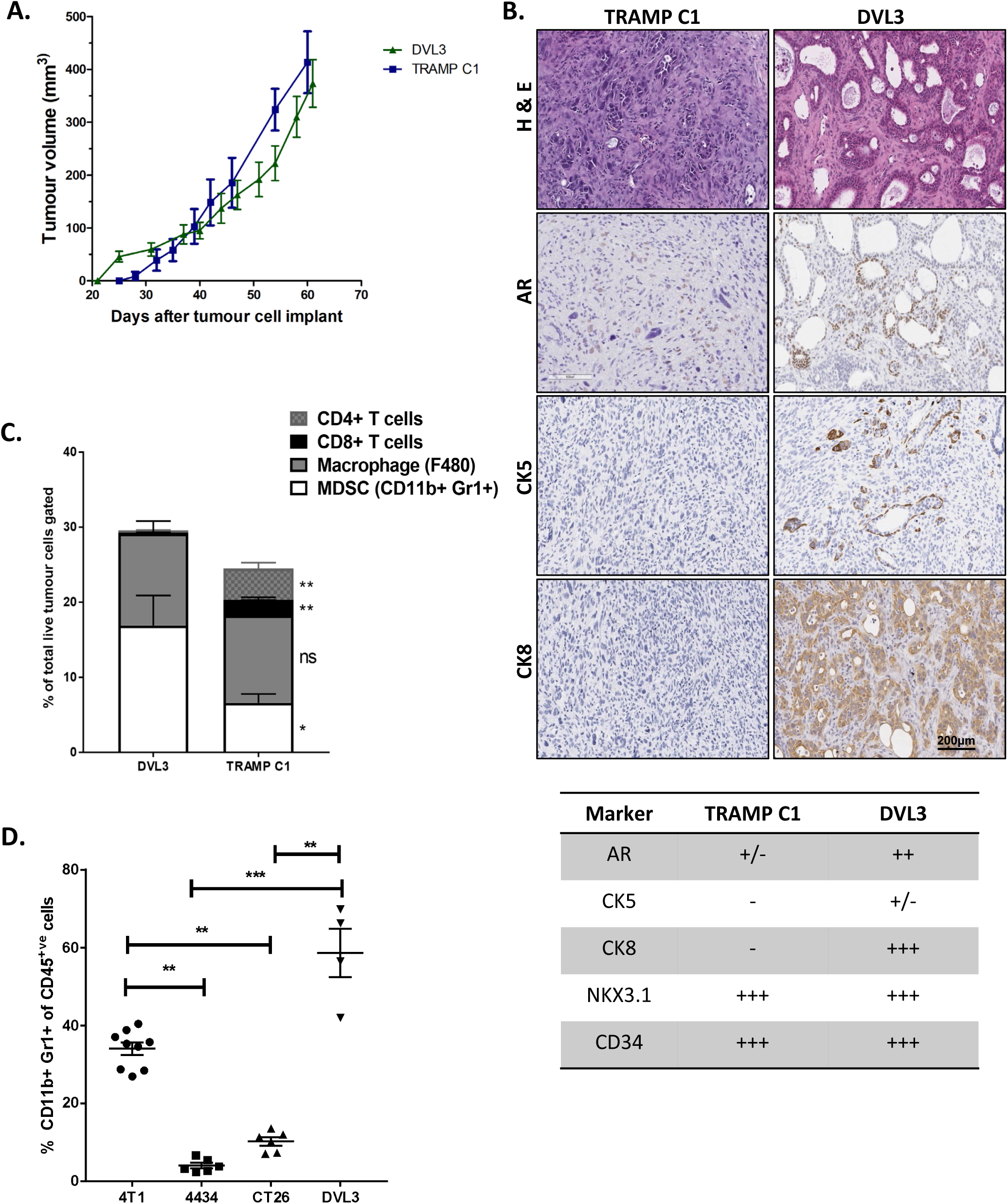
DVL3 prostate tumours replicate patient disease. **(A)** Tumour growth was comparable between TRAMP-C1 and DVL3 models, with both models taking ∼4 weeks to start to generate substantial tumours**. (B)** DVL3 tumours develop heterogeneous glandular morphology graded at Gleason 7, whereas TRAMP C1 tumours were undifferentiated with neuroendocrine features (H&E). DVL3 also expressed clinical prostate cancer markers; androgen receptor (AR), cytokeratin 5 (CK5) and cytokeratin 8 (CK8). TRAMP C1 expressed no CK8 or CK5 and less AR than DVL3 tumours. Histological evaluation of prostate specific markers in DVL3 and TRAMP C1 prostate tumours (- indicates absence, +/- heterogenous expression, and +s indicate the level of expression). Both DVL3 and TRAMP C1 express NKX3.1 and CD34 (Representative images in supplementary). **(C)** Flow cytometry in the DVL3 tumours compared to the TRAMP-C1 model demonstrates the proportion of helper T-cells (CD4+)and cytotoxic (CD8+) T-cells is significantly less in the DVL3 compared to TRAMP-C1 tumours. In addition the DVL3 presented with a significantly higher proportion of (CD11b+/Gr1+) positive cells compared to the TRAMP C1 tumours. Data represents mean ± SEM of at least (n=3) mice. **(D)** The DVL3 tumours also have higher baseline MDSCs, compared to other well characterised syngeneic murine models. Data represents mean ±SEM for at least (n=4) mice per group. * denotes p≤0.05, ** denotes p≤0.001 as assessed by unpaired t-test between each group.

DVL3 tumours displayed heterogeneous pathology with neoplastic, glandular structures akin to human acinar adenocarcinoma (**Supplemental Figure-3B**). Some regions of adenosarcoma were observed in the larger tumours as previously reported arising from trp53^-/-^/Pten^-/-^ Pb-Cre4 mice ^7^. In contrast, TRAMP-C1 tumours were uniformly undifferentiated and lacked any trace of glandular morphology **(Figure-2B).** Immunohistochemical staining revealed that the DVL3 tumours highly expressed CK8 throughout the entire tumour, particularly in the cells surrounding the lumen in the glandular structures **(Figure-2B)**. DVL3 tumours also had regions of CK5 positivity, with varied expression between the tumours. Conversely, in line with *in vitro* expression, TRAMP-C1 tumours lacked both CK8 and CK5. AR was present in the nucleus of cells throughout DVL3 tumours but was most apparent in clustered regions lining the lumen. TRAMP-C1 tumours had notably less AR staining and indeed some tumours lacked AR entirely **(Figure-2B)**. The clustered regions of AR positivity noted in the DVL3 tumours were not present in TRAMP-C1 tumours. To investigate if AR within the tumours was active, a downstream target of the AR; NKX3.1, was evaluated. NKX3.1 is a prostate-specific protein which is lost as PCa becomes castrate resistant ^18^. Both the DVL3 and the TRAMP-C1 tumours were highly positive for NKX3.1 and expressed across the tumour section **(Supplementary Figure-3A)**. Vascularisation of the models was also investigated via CD34 immunostaining, which highlights endothelial cells, no notable differences were observed between the models **(Supplemental Figure-3A).**

### DVL3 tumours have immunosuppressive microenvironment similar to human prostate adenocarcinomas

To assess the tumour immune microenvironment, flow cytometric analysis was performed for T-cells (CD4+, CD8+), macrophages (F4/80), and MDSC, identified as dual Gr1+/CD11+ cells. The DVL3 tumours have a significantly higher proportion of Gr1+/CD11b+ cells compared to the TRAMP-C1 tumour (16.90% vs 6.61%, p=0.04, **Figure-2C**). In contrast, both tumour lines have comparable level of macrophages (F4/80+) cells. The DVL3 tumours have significantly lower proportion of both cytotoxic CD8+ T-cells (0.187% vs. 2.05%) and CD4+ T-cells (0.36% vs. 4.31%) compared to the TRAMP-C1 tumours (p<0.005), similar to the immuno-suppressive microenvironment observed in PCa patients (11). MDSCs drive immune-suppression and can confer resistance to radiotherapy ^19^. Interestingly, the proportion of MDSCs in the DVL3 tumours is significantly higher compared to other syngeneic mouse models, such as CT26 (colorectal) and 4434 (melanoma) models, which are more sensitive to fractionated radiotherapy as demonstrated previously ^20^ (p<0.005, **Figure-2D**).

### Fractionated radiotherapy leads to marginal growth delay and alters the local tumour immune microenvironment

RT is a radical primary treatment administered to patients with high-risk localised PCa and also to patients with metastatic PCa. Therefore, therapeutic response to RT was evaluated in the DVL3 model. Established tumours (100mm^3^) were assigned to treatment groups (∼4 weeks post engraftment). The mice received clinically relevant single high dose of 8Gy or fractionated radiotherapy (2Gy per fraction administered over 5 consecutive days) (**Figure-3A**). DVL3 tumours responded to fractioned RT and exhibited a significant growth delay compared to non-treated controls (p=0.02, **Figure-3B**). Although there was no statistical difference in the size of tumours receiving a single high dose of 8Gy, both fractionated (5×2Gy) and single dose RT improved survival of mice carrying DVL3 allografts (p=0.03 and 0.02 respectively, **Figure-3B**). Conversely, neither the TRAMP-C1 tumour size nor survival was significantly affected by either RT regime (**Figure-3C**). This demonstrates DVL3 allograft tumours are more responsive to RT, in particular fractionated RT, compared to TRAMP-C1 which are known to be radioresistant ^21^.

**Figure 3:**
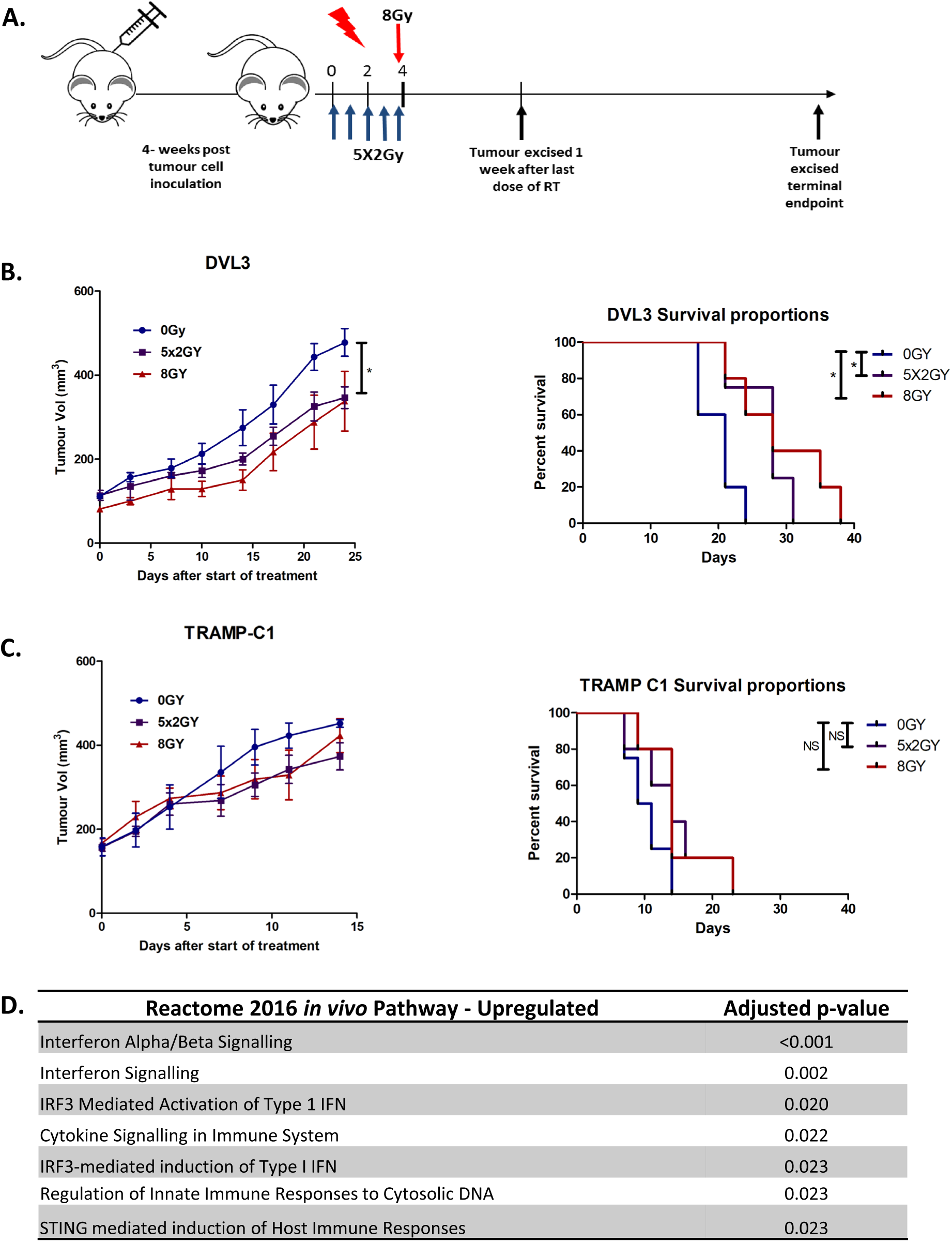
Radiotherapy resulted in tumour growth delay and differentially upregulated genes associated with STING activation and type-1 interferon signalling. **(A)** DVL3 or TRAMP C1 cells were engrafted into immune competent mice and allowed to reach a tumour volume of 100mm^3^. Mice were randomly assigned to no treatment (0Gy), fractionated dose (5×2Gy) or single high dose (8Gy) RT group. **(B)** DVL3 tumours were sensitive to fractionated RT but no significant difference was seen with single high dose RT. Kaplan Meier curves showing survival in mice engrafted with DVL3 tumour cells was significantly improved for both treatment regimens (5×2Gy and 8Gy) compared to for mice treated with 0Gy (5×2Gy p=0.0255 and 8Gy p=0.0386) **(C)** TRAMP C1 tumours did not significantly respond to radiotherapy. There was no change in survival with radiation in the mice engrafted with TRAMP C1 cells. **(D)** Pathways enrichment was preformed using the online software Enricher for differential gene expression comparing the fractionated (5×2Gy) tumours to non-irradiated control *in vivo*. Upregulated pathways enrichment in the Reactome 2016 for transcripts with a significance of greater than 0.05 and fold increase of ≥1 (Inert statistical test you used for this).

RT is known to induce immunogenic changes in tumour cells and can re-calibrate the immune contexture of the tumour microenvironment ^22^. Therefore, further investigation via RNA-Seq was performed on DVL3 tumours treated with fractionated radiation and un-irradiated control tumours. Interestingly, ENRICHR pathway analysis on differentially expressed genes highlighted a significant enrichment for pathways involved in interferon signalling (p<0.001) and STING activation (p<0.023) up-regulated in the fractionated tumours (**Figure-3D**).

STING dependent cytosolic DNA sensing promotes type-1 interferon response and is critical for activation of innate immunity (19). GSEA using gene sets for MDSCs and macrophages revealed significant up-regulation of the gene signatures; however; no increase was observed for genes associated with T-cells (**Figure-4 A-C**). In order to establish phenotypic significance of the activation of type-1 IFN-pathway, the expression of STING was evaluated via immunohistochemistry. Fractionated RT led to an abundant increase in STING expression, primarily from CK8+ tumour cells (**Supplementary Figure-3C**). The increase in expression of STING on tumour cells correlated with a marginal increase in myeloid cell infiltrates (CD11b+) in the irradiated tumour (15.2% versus 23.3%, p=0.08). However, RT did not result in a significant increase in either CD8+ or CD4+ T-cells (**Figures-4B and C**). A marginal increase in NK cells (Nkp46) was observed which were predominantly localised around the necrotic areas in tumours **(Figure-4B and C**). In addition, macrophage infiltration showed no increase as measured using (F4/80+) cells in the irradiated DVL3 tumours (**Figure-4B and C**). Taken together with the GSEA, our results suggest that the activation of type-1 signalling in the irradiated DVL3 tumours could be due to influx of MDSCs and activation on innate immunity.

**Figure 4:**
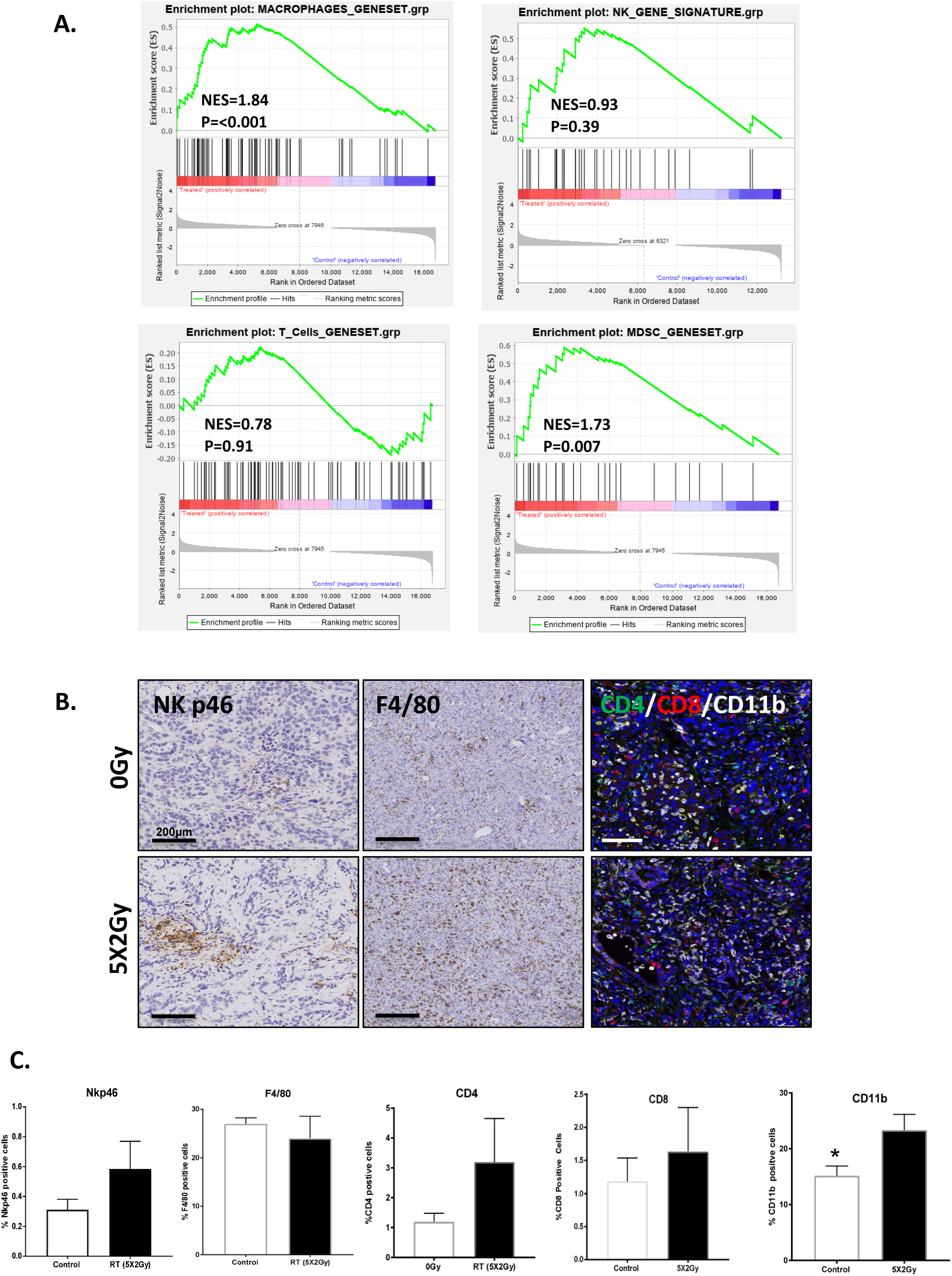
Fractionated radiotherapy lead to significant upregulation of gene signatures associated with MDSC and macrophages, which correlated with an increase in infiltration of myeloid cells in the irradiated tumours. GSEA enrichment analysis of fractionated (5X2Gy) treated tumours demonstrates an upregulation of **(A)** macrophage and MDSC signatures, but no enrichment of NK cell or T-cell signatures when compared with non-irradiated tumours. **(B)** Representative chromogen staining for macrophages (F/480), natural killer (Nkp46) and multiplex staining images for (CD4 (green), CD8 (red) and CD11b (white)), in the DVL3 tumours**. (C)**. Quantification for macrophages (F4/80) in stained sections. Quantification for NK-cells in tumours showing a marginal increase in irradiated tumour. Quantification for myeloid (CD11b+ cells); cytotoxic (CD8+), and helper (CD4+) T-cells. Data represents mean ± SEM percentage positive in at least (n=3) mice per treatment group.

## Discussion

Whilst significant progress has been made in the treatment of metastatic, castrate-resistant prostate cancer (CRPC), the majority of these cancers are incurable. In order to have greater impact, it is important to understand how PCa evolves from localised disease so that more effective treatment combinations can be developed. The narrow range of pre-clinical models limits therapeutic development in prostate cancer. This study has developed two novel murine cell models; mPECs, which model normal prostate epithelium and may prove useful to engineer and test the impact of other clinically relevant driver mutations, and the tumorigenic DVL3 cell line. DVL3s were generated through deletion of genes that are frequently mutated in human PCa and are implicated in aggressive forms of the disease ^7^; thus, avoiding the introduction of tumorigenic viral proteins such as SV40 large T-antigen, which was used to generate the TRAMP-C1 cell line ^6^. Although the TRAMP-C1 cell line has contributed significantly to PCa research, recent evidence indicates it generates neuroendocrine tumours, which clinically equates to only 0.5–2% of all PCa cases ^23^. Conversely, the DVL3 develop tumours that represent an adenocarcinoma phenotype much more akin to most human PCa.

Transgenic models of PCa, and subsequent cell lines, generated from the trp53^-/-^/Pten^-/-^ model form heterogeneous tumours **(Figure-2B)** (13). Arguably, heterogeneity is one of the greatest benefits of this model, as most cell-based models are thought to be clonally selected due to genetic drift. As PCa is a multifocal heterogeneous disease the derivation of the bulk DVL3 population allows for research into PCa evolution in responses to treatment ^24^. Tumours formed by the DVL3s maintained the heterogeneous CK8+/CK5+ glandular structures and regions of AR positivity, similar to the tumours from which they were derived. Additionally, unlike other trp53^-/-^/Pten^-/-^ murine cell lines, previously described in the literature, the DVL3 can be allografted, presenting with immunological cold tumour immune microenvironment, mirroring observations seen clinically ^7,25^.

Development of immune therapies as a method of harnessing the immune system’s anti-tumour effects has recently gained traction ^25^. Immune checkpoint inhibitors (ICI) have been used successfully to treat melanoma and bladder cancers; however, their impact on PCa is more limited ^24^. Unlike melanoma and lung cancer, PCa has low levels of infiltrating CD8+ T cells ^24,25^. The immune microenvironment of PCa has been implicated in a number of other ways including: host physiological factors, DNA damage response defects, low levels of tumour-associated antigens and the association of these factors with PTEN loss ^24^. PTEN loss occurs in 20% of primary PCa patients and has been associated with poorer overall survival ^7,24^. Additionally, melanoma patients with PTEN loss also show lower levels of tumour infiltrating lymphocytes (TIL) ^24^. We have confirmed that this is also true in the DVL3 allografts, underscoring the more immuno-suppressed microenvironment of these tumours compared to the TRAMP-C1 tumours **(Figure-2D)**.

Comparing the DVL3 allografts to published transgenic models driven by Pten-deletion highlights other similarities and opportunities for future studies. For example Pten-deletion is known to lead to the expansion and immunosuppressive activities of Gr1^+^/CD11b^+^ myeloid derived suppressor cells (MDSC) ^26^. High numbers of MDSCs in a Pten^-/-^ setting provide a protective effect on PCa cells from senescence, therefore sustaining tumour growth ^26,27^. Mechanistically this has been observed to occur in CRPC patients and in transgenic prostate cancer models through the activation of the AR signalling due to paracrine IL-23 secretion by MDSCs ^28^. Furthermore, influx of tumour MDSCs has been detected following RT, an observation which has been previously associated with the RT-induced up-regulation of STING expression in tumour cells ^19,29,30^. Targeting MDSC, through IL-23 inhibition, in combination with RT and ADT could therefore provide a survival benefit for PCa patients. The DVL3 model recapitulates clinically relevant immune microenvironment features that have been previously observed in PCa transgenic models following RT, such as influx of MDSC **(Figure-4)** ^19,26,28^.

The DVL3 allograft model represents a significant improvement in prostate cancer research. DVL3 tumours more accurately recapitulate patient disease molecularly, histologically, and morphologically compared to alternative allograft models. Furthermore, they can be generated quickly and inexpensively compared to standard transgenic models, and tumours are readily accessible for therapeutic intervention. Importantly, DVL3 cells can be syngeneically engrafted into immune component hosts and respond to standard of care RT, with a similar immune influx. These data highlight the DVL3 cells as an improved PCa model, which may be used for the development and validation of novel therapeutics for prostate cancer, in particular immune-modulatory agents in combination with RT.

## Additional Information

### Ethics approval and consent to participate

All procedures were performed in accordance with the Animal Scientific Procedures Act of 1986 (UK) (Project Licence Number PL2775 and PCC943F76) which was issued by the home office. Protocols were approved by the animal welfare and ethical review body at both Queen’s University Belfast and the University of Manchester. Animals were housed in individually ventilated cages on a 12:12 light:dark cycle and ad libitum access to food and filtered water. Prior to each in vivo experiment, cells were screened for mycoplasma contamination.

### Consent for publication

N/A

### Data availability

RNA-seq files are currently being deposited on the Gene Expression Omnibus (GEO) database. GSE number – Pending

### Conflict of interest

The authors declare no potential conflicts of interest.

### Funding

CH was funded by the Gracey Foundation. DM, RES, LDP, AP, SJ, SMD, TI, and IGM were supported by the Belfast-Manchester Movember Centre of Excellence (MA-CE018-002), funded in partnership with Prostate Cancer UK. SLE, PBM and RW were funded by Prostate Cancer UK (PCUK PG13-021). NB funded by Breast Cancer Now (2012MAYSF122). TI supported by NIHR Manchester Biomedical Research Centre and CRUK programme grant. JH funded by Cancer Research UK Programme grant (A17737)

### Authors’ contributions

CMH, DM, SLE - designed experiments and helped write the manuscript.

RES, AM, LDP, AP, PO, NEB, JH, SMD, IGM - Helped design experiments and gave expert advice.

IGM, SJ, PBM, RW, SMD, TI - secured funding for individuals and consumables.

## Acknowledgements

We acknowledge the generous gift of mouse prostate tissue from the David Waugh laboratory from which the mPEC and DVL3 cell lines were generated

