## Supplementary figures and tables for "Development of a novel allograft model of prostate cancer: a new tool to inform clinical translation"

### Supplementary information

1. Supplementary Figure Legends
2. Supplementary Table Legends
3. Supplementary Methods

### 1. Supplementary Figures

**Supplementary Figure-1: Methodology for the isolation and culturing of murine primary prostate cells lines.**

**A)** Schematic of the protocol used to digest and isolate the DVL3 and mPECs primary murine cell lines from murine prostate tissue. **(B)** DVL3 and TRAMP C1 cells were treated with enzalutamide72hr after treatment plates were treated with Alamar Blue and absorbance measured (n=5). Cell viability was measured and made relative to DMSO control. IC50 values demonstrating the DVL3 were more sensitive than the TRAMP C1. Error bars represents the ± SEM of independent experiments. Unpaired t test statistical analysis; * (p=<0.05).

**Supplementary Figure-2: Gating strategy to identify MDSCs type population in the dissociated DVL3 tumours.**

(**A**) Schematic outlining the gating strategy to identify Gr1/ CD11b+ positive cells of total CD45 cells from the DVL3 and TRAMP-C1 tumours. **(B)** Individual flow cytometry data comparing the levels of Gr1+/ CD11b+, F4/80+, CD8+ and CD4+ cells in DVL3 tumours and TRAMP-C1 tumours for Figure-2D.

**Supplementary Figure-3: Additional characterisation of DVL3 allografts**

**(A)** Representative images demonstrating expressing of NKX3.1 and CD34 in both DVL3 and TRAMP C1. Histological evaluation of these markers are reported in **Figure-2A.** **(B)** Representative images of the DVL3 tumours demonstrating glandular structures akin to human acinar adenocarcinoma. **(C)** Representative multiplex immune-histochemical staining for STING (Green) and CK8 (Red) in the DVL3 tumours treated with fractionated RT or non-treated control. RT led to an increase expression of STING in the tumour cells as demonstrated from double staining for STING with CK8 positive. **(D)** Mean survival of mice, in days, after the final administration of RT. Data represents mean ± SEM percentage positive in at least (n=3) mice per treatment group. Unpaired t-test statistical analysis; * ($p<0.05$). Pathways enrichment was preformed using the online software Enricher for differential gene expression comparing the fractionated (5x2Gy) tumours to non-irradiated control in vivo. Downregulated pathways enrichment in the Reactome 2016 for transcripts with a significance of greater than 0.05 and fold decrease of ≥-1.

### 2. Supplementary Tables

**Supplementary Table-1:** List of qRT-PCR primers.

**Supplementary Table-2:** List of antibodies.

### 3. Supplemental Methods

**Cell Line Derivation**

Tissue fragments were centrifuged at 2000 rpm to remove extraneous blood and fat, PBS was removed, and remaining tissue digested in 1mg/ml collagenase/dispase in PBS for 30min at 37°C. Tissue fragments were further manually dissociated and the supernatant containing prostate cells collected. Collagnease/dispase was inactivated with EDTA at this time. The digestion process was repeated twice with remaining tissue fragments. Supernatant from the digestions was pooled and centrifuged at 2000 rpm and prostate cells were re-suspended in RPMI supplemented with 10% FBS, 1X pen/strep, 100nM DHT and rock inhibitor. These cells were then incubated for 10 minutes at 37°C to allow for prostate fibroblasts to adhere to the tissue culture plastic. Remaining cell suspension, enriched for prostate epithelium was transferred to a new culture flask. This process of differential plating, allowing for initial adherence of fibroblasts and removal and retention of less adherent epithelial cells, was repeated for 10 passages, at which time the rock inhibitor was removed and the population remained homogenous.

***Cell Line Generation and Maintenance***

Mouse prostate epithelial cells (MPEC) were generated from dorsal, ventral and lateral lobes of the prostate from Probasin Cre$^{-/-}$ (Pb-Cre4) mice. Murine prostate cancer cells (DVL3) were generated from tumours derived from the dorsal, ventral and lateral prostate lobes of a trp53$^{-/-}$/Pten$^{-/-}$ Pb-Cre4 mouse. Tissue was manually dissociated under sterile conditions and cell lines were generated as fully described in **Supplemental Methods and Supplementary Figure-1A**.

The TRAMP-C1 murine prostate carcinoma cells were purchased from ATCC and maintained in DMEM high glucose medium, supplemented with 4mM L-Glutamine, 5% FBS, 5% Nu Serum, 0.005mg/ml of Bovine insulin, and 10nM Dehydroisoandrosterone (Sigma). The DVL3 cell lines were maintained in RPMI-media supplemented with 10% FBS, L-Glutamine and 100nM DHT.

***RNA Isolation and cDNA synthesis***

Total RNA from cells was isolated using TriPure Isolation reagent (Roche) and phenol-chloroform extraction and cDNA synthesised using first strand cDNA (Roche) according to the respective manufacturer's instructions.

***qRT-PCR***

mRNA levels were determined by RT-PCR and measured on Roche LighCycler®480 II. All primers were obtained from Eurofins and sequences are detailed in (**Supplemental Table-1**) mRNA expression was quantified relative to house-keeping genes; B-actin.

***Western blotting***

Cells were lysed with Erythrocyte Lysis Buffer (0.5mM DTT, 5mM EDTA (pH 8), 50mM HEPES (pH7.5), 0.1% IGEPAL, 250mM NaCl) supplemented with Protease Inhibitor Cocktail (Roche), centrifuged and supernatant containing protein analysed by SDS Page (Invitrogen). Proteins were transferred onto a nitrocellulose membrane, blocked for 1 hour in 5% milk TBS-T prior to incubation with primary antibody at 4°C overnight, concentrations are detailed in (**Supplementary Table-2**). Complementary secondary antibodies were used at 1:3000. Membranes were visualised on a Syngene G:Box imager using reagent (Merck).

***Allograft modelling***

C57BL/6 male mice (8 weeks old) were obtained from Harlan, UK. Animal experiments were approved by a local ethical committee and performed under a United Kingdom Home Office project licence. Prior each *in vivo* experiment, cells were screened for mycoplasma contamination and MHV. Mice were housed on a 12/12 light/dark cycle and were given filtered water and fed ad libitum. Mice were inoculated subcutaneously with either $5x10^6$ TRAMP-C1, $1x10^6$ DVL3 cells or $1x10^6$ mPEC cells.
